# The central autonomic system revisited – convergent evidence for a regulatory role of the insular and midcingulate cortex from neuroimaging meta-analyses

**DOI:** 10.1101/2022.05.25.493371

**Authors:** Stefania Ferraro, Benjamin Klugah-Brown, Christopher R Tench, Mercy Chepngetich Bore, Anna Nigri, Greta Demichelis, Maria Grazia Bruzzone, Sara Palermo, Weihua Zhao, Shuxia Yao, Xi Jiang, Keith M Kendrick, Benjamin Becker

## Abstract

The autonomic nervous system regulates dynamic body adaptations to internal and external environment changes. Capitalizing on two different algorithms (Analysis of Brain Coordinates and GingerALE) that differ in empirical assumptions, we scrutinized the meta-analytic convergence of human neuroimaging studies investigating the neural basis of peripheral autonomic signal processing. Among the selected studies, we identified 42 records reporting 44 different experiments and testing 792 healthy individuals.

The results of the two different algorithms converge in identifying the bilateral dorsal anterior insula and midcingulate cortex as the critical areas of the central autonomic system (CAN). However, whereas the bilateral dorsal anterior insula appears to be involved in processing autonomic nervous system signals regardless of task type, activity in the midcingulate cortex appears to be primarily engaged in processing autonomic signals during cognitive tasks and task-free conditions. Applying an unbiased approach, we were able to identify a single functionally condition-independent circuit that supports CAN activity. Although partially overlapping with the salience network, this functional circuit includes, in addition to the bilateral insular cortex and midcingulate cortex, the bilateral inferior parietal lobules and small clusters in the bilateral middle frontal gyrus. Our results do not support the hypothesis of divergent pathways for the sympathetic and parasympathetic systems or a robust involvement of the default mode network, particularly during parasympathetic activity. However, these results may be due to the relatively low number of studies investigating the parasympathetic system (12%), making our results more consistent with the central processing network of sympathetic activity.

Remarkably, the critical regions of the CAN observed in this meta-analysis are among the most reported co-activated areas in neuroimaging studies and have been repeatedly shown as being dysregulated across different mental and neurological disorders. This suggests that the central dynamic interaction maintaining bodily homeostasis reported in several brain imaging studies may be associated with increased autonomic nervous system engagement and that disruptions in this interplay may underpin unspecific pathological symptoms across mental and neurological disorders.

## 1. Introduction

The autonomic nervous system (ANS) represents an integral part of the peripheral nervous system and serves an essential role in maintaining homeostasis by regulating involuntary physiological functions, including regulation of the cardiovascular, respiratory, digestive, and genitourinary systems as well as exocrine and endocrine glands. The ANS supports these fundamental functions with dynamic adaptations of the body in response to changes in the internal and external environment (Cannon, 1929), based on ongoing or planned behavior, through bidirectional innervations (afferent and efferent) between the brain and target organs (i.e., heart, smooth muscle, viscera, immune tissue, and exocrine and endocrine glands). This complex system consists of three components (Cardinali, 2017): the parasympathetic system, regulating internal bodily functions (i.e., “rest and digest state”), the sympathetic system, orchestrating physiological responses in response to arousing stimuli (e.g., “fight-or-fly responses”), and the enteric system, which can operate autonomously from the brain and spinal cord (Gershon & Nakamura, 2019) but will not be further considered in the present work. In a continuous bidirectional exchange of information and fine-grained interactions between the periphery and the brain afferent signals from the ANS (through viscerosensory arms) reach the “central autonomic system” (CAN) of the brain, thus representing and modulating interoceptive information in the brain, while efferent signals from the CAN (through motor arms) reach target organs and modulate bodily functions (Cardinali, 2017).

The dynamic bidirectional interplay between internal body states and their central interoceptive representation shapes several mental processes (Critchley & Harrison, 2013). The close link between ANS signals and mental processes has been hypothesized since the late 19th century by the James-Lange theory (W. James, 1994; Lange & Haupt, 1922) proposing that peripheral ANS signals represent precursors of emotions. Subsequent studies have found some evidence for this hypothesis by identifying emotion-specific fingerprints of ANS signals (Collet et al., 1997; Ekman et al., 1983; Harrison et al., 2010). More recent evolutions of this conceptualization, such as the “somatic marker model” (Damasio, 2003) or “interoceptive-based model” (Critchley & Harrison, 2013) hypothesize that ANS signal fingerprints additionally modulate cognitive processes such as attention, decision-making, and memory (Critchley et al., 2013).

Animal studies have mapped the neurobiological systems involved in CAN in great detail. In particular brainstem systems, where the parabrachial and tractus solitarius nuclei constitute the main inflow and outflow structures for autonomic signals, have been mapped. These nuclei receive parasympathetic (via cranial and sacral neural pathways) and sympathetic (via thoracolumbar neural pathways) afferent inputs and transmit the processed information to the hypothalamus, amygdala, thalamus, and cortical regions, with key projections into the insular and medial prefrontal cortex. The hypothalamus (lateral and paraventricular nucleus) and brainstem nuclei (periaqueductal gray matter, reticular formation, in addition to the aforementioned tractus solitarius and parabrachial nucleus) constitute sympathetic and parasympathetic efferent nodes, and are regulated by the amygdala, insula, and prefrontal cortex (Saper & Stornetta, 2015).

Over the past two decades, an increasing number of neuroimaging and neurostimulation approaches have been used aimed at determining systems that mediate peripheral sympathetic and parasympathetic activity in humans. These studies have commonly employed a variety of ANS measures and modulations (e.g., heart rate variability, skin conductance, muscle and skin sympathetic nerve activity, skin temperature, vagus nerve stimulation) during different mental processes (i.e., cognitive, affective, somatosensory-motor, at rest) and broadly confirmed a crucial role of insular and frontal midline regions in regulating autonomic activity (Beissner et al., 2013). However, despite an increasing number of studies the functional architecture and dynamic system-level organization of the ANS in humans – especially at the forebrain level which is difficult to determine in translational animal models - remains debated, e.g., with respect to hemispheric lateralization, large-scale dynamic networks, and domain-general versus-specific engagement in affective and cognitive processes.

A general and debated question relates to the hemispheric organization of the ANS, with previous studies suggesting lateralization to the right hemisphere for the sympathetic system and lateralization to the left hemisphere for the parasympathetic system (A D Craig, 2005; Oppenheimer & Cechetto, 2016; Thayer & Lane, 2009). More recently, it has been suggested that ANS activity is regulated by lateralized circuits belonging to the salience network, with the left ventral anterior insula cortex modulating the parasympathetic system while the right hypothalamus/amygdala modulates the sympathetic system (Sturm et al., 2018). However, despite accumulating evidence, the last comprehensive meta-analysis of neuroimaging studies on the human CAN (Beissner et al., 2013) did not confirm hemispheric lateralization. Similarly, it remains unclear whether the sympathetic and parasympathetic systems map onto divergent large-scale networks. The early neuroimaging meta-analysis (Beissner et al., 2013) suggests divergent networks for the two ANS systems, e.g., reporting that sympathetic activity would map onto key regions of the executive control and salience network, whereas parasympathetic activity would map onto the default mode network (DMN), with only the amygdala and right anterior insular cortex involved in processing both sympathetic and parasympathetic signals (Beissner et al., 2013). The salience network (Seeley et al., 2007b) receives continuous interoceptive information and triggers visceromotor changes to match current or anticipated metabolic demands and has been identified as a regulator of the ANS (Beissner et al., 2013; Benarroch, 1993; Critchley & Harrison, 2013; Saper, 2002; Sturm et al., 2018). In contrast, previous studies revealed inconsistent results regarding the role of the DMN in maintaining homeostasis (Dhond et al., 2008; Nagai et al., 2004; Wong et al., 2007). Finally, despite the long-standing hypothesis on the influence of peripheral ANS signaling on cognitive and emotional processes, it has not been established whether representations of central ANS activity differ depending on the domain involved. The previous meta-analysis (Beissner et al., 2013) examined affective, somatosensory-motor, and cognitive paradigms and identified both shared (right anterior and posterior insula cortex, left amygdala, and midcingulate cortex) and domain-specific representations, such that cognitive tasks primarily recruited regions involved in sympathetic modulation whereas affective and somatosensory-motor tasks engaged regions involved in both sympathetic and parasympathetic modulation.

Despite the important contributions of the seminal meta-analysis of functional neuroimaging of the human ANS by Beissner et al. (Beissner et al., 2013), the number of original studies has greatly increased since its publication nearly 10 years ago and it is now an opportune time to revisit the central organization of the ANS in humans by capitalizing not only on the increased number of original studies but also methodological and conceptual advances. For instance, numerous studies have combined experimental progress with resting-state functional magnetic resonance imaging (fMRI) assessments or vagus nerve stimulation over the last years and methodological approaches for neuroimaging metaanalyses have improved considerably. For example, problems in the early implementation of coordinate-based meta-analyses (e.g., in GingerALE) have been addressed (Eickhoff et al., 2009, 2012, 2017) and novel model-based approaches developed allowing meta-analytic examination with less empirical assumptions (C R Tench et al., 2022) facilitating more robust results. Moreover, CAN dysregulations in clinical contexts have received growing attention (Johnson & Wilson, 2018).

Against this background, the present pre-registered meta-analysis capitalizes on the increasing number of original neuroimaging studies examining CAN in humans and on the advancements in meta-analytic methods, including functional and network-level characterization to reach two key goals. First, to identify brain systems that are robustly involved in the central processing of ANS signals (corresponding to the CAN), and second, to define whether neural representations of the CAN vary according to the affective functional domain (affective, cognitive, somatosensory-motor, task-free experiments), or according to the autonomic system involved (sympathetic and parasympathetic). With respect to the latter goal, we examined the impact of negative affective load by comparing studies employing affective and somatosensory-motor approaches (mainly using acute pain induction) with studies employing cognitive and task-free approaches that lack a strong negative affective component.

Moreover, to answer whether CAN areas vary according to the ANS system involved (sympathetic and parasympathetic system), we defined the physiological measures taken in the original studies which identify the activity of a specific ANS system - (e.g., high-frequency heart rate variability is an index of the parasympathetic activity or muscle sympathetic nerve activity is an index of sympathetic activity) – or which are indices of a combination of the activity of both systems (e.g., pupillary diameter). We validated the robustness of our results using two meta-analytic approaches with different coordinate-based meta-analytic (CBMA) empirical assumptions, namely GingerALE (Eickhoff et al., 2009, 2012, 2017), the most popular algorithm of performing CBMA, and the novel developed Analysis of Brain Coordinates (ABC) (C R Tench et al., 2022). GingerALE requires the choice of several empirical assumptions that, in principle, can influence the results (Ferreira & Busatto, 2010), while ABC is a novel and validated method that requires only minimal empirical assumptions, namely grey-matter, whitematter (WM), or whole-brain (WB) volume being the statistical thresholding conceptualized as the minimum replicates considered adequate by the experimenter. After determining general and domain-specific CAN systems, we employed further meta-analytic strategies to promote a functional characterization of the identified regions at the network level and thus to determine which large-scale networks are engaged in regulating ANS, in particular the salience and default mode networks (DMN).

## 2. Materials and Methods

### 2.1 Study selection

This meta-analysis was pre-registered on PROSPERO (registration number: CRD42021270736) and followed the PRISMA reporting guidelines (Moher et al., 2009) (figure 1). Public databases of biomedical and life science literature reports, namely Pubmed, Web of Sciences, Scopus, and the Neurosynth database were searched for functional magnetic resonance imaging (fMRI) or positron emission tomography (PET) studies investigating CAN activity in healthy adult populations during task-based and task-free conditions. The following keywords were used to search for papers present in the databases from January 1995 to 10th of August 2021: ((“autonomic” OR “sympathetic” OR “parasympathetic” OR “vagal” OR “vagus”) AND (“fMRI” OR “functional magnetic resonance” OR “functional” OR “brain activation” OR “neural activity” OR “BOLD” OR “PET”) AND (“heart rate” OR “respiration” OR “skin conductance response” OR “skin response” OR “pupil” OR “skin temperature” OR “blood pressure” OR “electrogastrography”)). We performed another search on 7th of March 2022 to determine newly published papers before the final submission. For each identified paper, we collected the title, author names, and date of publication. We merged the results from the different databases and identified and removed all duplicates. A unique identification number was assigned to the remaining papers. Two independent reviewers (S.F. and G.D) excluded records not relevant for the current meta-analysis based on the title and abstracts. Records were considered excluded at this step when they were excluded by both reviewers. The remaining studies were then reviewed by means of a careful reading of the full text to identify the investigations fulfilling the criteria of inclusion (see figure 1). Studies were included if they: 1) investigated CAN activity in healthy individuals using fMRI or PET; 2) recorded ANS activity or employed a direct stimulation of the ANS (e.g. transcranial vagus nerve stimulation) during neuroimaging; 3) employed the recorded ANS activity in the analyses of the neuroimaging data as a variable of interest (not as nuisance or noise correction variable); 4) employed random effect (Friston et al., 1999) or multivoxel pattern analysis at a whole-brain level (studies employing only regions of interest analyses were excluded) in a sample of at least 10 healthy individuals; 5) reported results of the experimental conditions investigating the CAN at a whole-brain level as coordinates in the standard Talairach & Tournoux (TAL) (Talairach, 1988) or Montreal Neurological Institute (MNI) (Evans et al., 1993) stereotactic space; 6) reported results as significant for voxel-wise corrected thresholds or voxel-wise uncorrected thresholds but corrected for cluster size or uncorrected thresholds of p < 0.001.

**Fig. 1.**
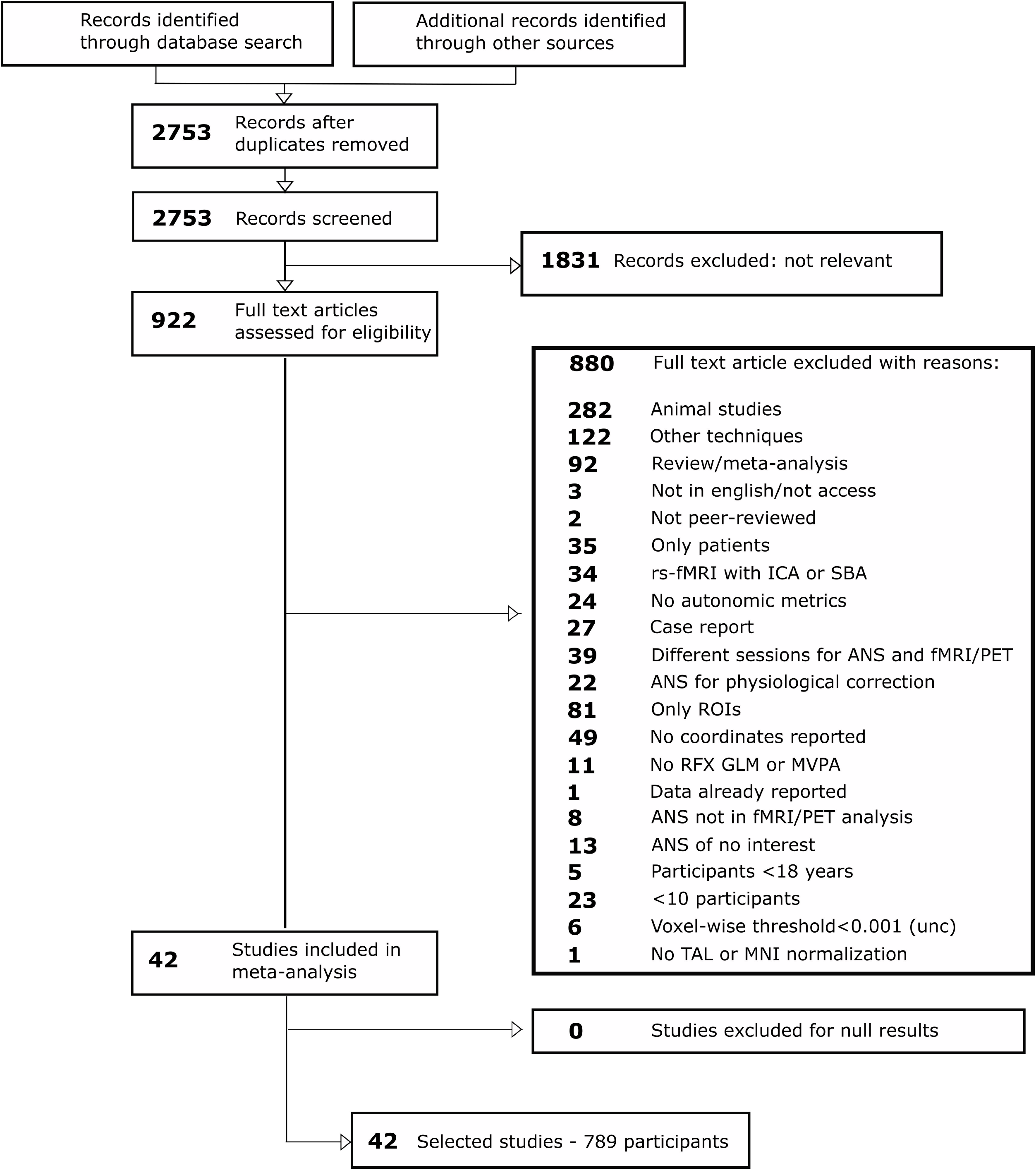
Flowchart of the screening process of the identified studies according to the PRISMA guideline.

### 2.2. Data extraction

For each selected record, one of the authors (S.F.) extracted the imaging technique (fMRI or PET), ANS measures employed in the analysis, sample size, coordinates and corresponding statistical values, and standard space (TAL or MNI). Another author (M.C.B.) independently verified all extracted data. When inconsistencies were observed, data were re-evaluated and corrected. An automatic diagnostic procedure was additionally employed on the extracted coordinates implemented in NeuRoi (Christopher R Tench et al., 2013). When potential errors were detected, coordinates were re-checked and corrected. All analyses were conducted only after performing these steps. Following the strategy in the previous meta-analysis on the topic by Beissner et al. (Beissner et al., 2013), the selected studies were classified according to two dimensions: the type of experiment (*task dimension*), used to study the ANS, and the investigated ANS system (*autonomic dimension*). With respect to *task dimension*, we classified the selected records as task-free (namely, resting-state studies) or task-based investigation. This latter category was further subdivided according to the investigated domain (affective, cognitive, or somatosensory-motor domain). Relative to the *autonomic dimension*, the selected studies were grouped in the following categories in agreement with the statements reported in the manuscripts: studies investigating the sympathetic, or the parasympathetic or the sympathetic/parasympathetic system activity. As explicitly mentioned in the study, we considered measures of the sympathetic nervous system as skin sympathetic nerve activity, muscle sympathetic nerve activity, skin conductance response, finger temperature or sympathetic stimulation with epinephrine, baroflex unloading or carotid sucks. We considered studies as investigating parasympathetic nervous system activity when they employed high-frequency heart rate variability and direct stimulation of the vagus nerve. We considered studies as investigating sympathetic/parasympathetic system activity when they employed different measures of heart rate variability (except high-frequency heart rate variability), pupil size changes, and tachigastria. The classification of the studies was performed by S.F. and then checked by an independent reviewer (A.N.).

### 2.3. Meta-analysis approaches: Analysis of Brain Coordinate and Ginger-ALE algorithms

CBMA algorithms test the null hypothesis that the peak coordinates reported in the selected studies are uniformly distributed throughout grey matter (Turkeltaub et al., 2002). However, different CBMA approaches rely on different empirical parameters and assumptions and thus can produce different results. To increase the robustness of our results we, therefore, employed two different tools: Analysis of Brain Coordinates (ABC) (C R Tench et al., 2022) and GingerALE (Eickhoff et al., 2009, 2012).

#### Analysis of Brain Coordinates (ABC)

There are multiple algorithms for performing CBMA, each with specific assumptions and each potentially producing different results (Ferreira & Busatto, 2010; C R Tench et al., 2022). ABC is a new model-based method with very few a priori assumptions, implemented in the NeuRoi image analysis software (https://www.nottingham.ac.uk/research/groups/clinicalneurology/neuroi.aspx}) (C R Tench et al., 2022). By removing the need for some empirical features commonly used in CBMA, the impact of the algorithm on the results is more transparent. The algorithm also makes the statistical thresholding more transparent by relating it to a minimum proportion of studies contributing to a valid clusters of coordinates, which may be easier to underatand than the typical thresholding schemes where the threshold relates to the rate of results expected under the null hypothesis. Moreover, ABC has the facility to include covariates that it incorporates into binary logistic regression models for each valid cluster of coordinates for further analysis. The default threshold of at least 10% of studies contributing coordinates to valid clusters.

#### GingerALE

The GingerALE algorithm(v. 3.0.2) performs a voxel-wise meta-analysis employing the number of participants of each selected study to smooth, with a Gaussian kernel, the identified foci and to produce Modelled Activation Maps (Eickhoff et al., 2009, 2012). The maps from the original studies are next merged in a single activation likelihood estimate (ALE) image to produce normalized histograms. By simulating random data a permutation test is used to define the statistical threshold to identify brain areas whose values are beyond chance level (Eickhoff et al., 2009, 2012). As recommended, we thresholded our ALE maps at cluster-level inference (cluster-level family-wise error p<0.05) and employed an uncorrected cluster forming threshold of p<0.001 (Eickhoff et al., 2012).

### 2.4. Meta-analyses

We assessed whether specific task-dimension experiments clustered in a specific autonomic-dimension by employing the Freeman–Halton extension of the Fisher exact probability test (three-by-four contingency table). Results (p = 0.13) indicated that task-dimension and autonomic-dimension experiments were independent. In accordance with the aims of the present study the following three meta-analyses were performed in TAL space (Talairach, 1988):

1. A pooled meta-analysis was used to determine brain systems robustly involved in CAN activity across all selected studies and independently from the two identified dimensions (namely, *task-dimension* and *autonomic-dimension*). This meta-analysis was conducted with both algorithms (i.e., ABC and GingerALE).
2. Task-dimension meta-analysis was conducted to investigate whether the task domains would impact the identified brain systems. To avoid power reduction in separate meta-analyses we capitalized on the covariate regression approach of ABC and included task-domain as covariate of interest. To this end the tasks were separated by their negative affective component, such that somatosensory-motor experiments (conducted mainly with pain and nausea inducing stimuli, or acupuncture) and affective experiments (conducted mainly with fearful or disgusting stimuli) inducing ANS reactivity by a strong negative affective stimulation were distinguished for task-free and cognitive experiments which usually lack these negative affective components.
3. Autonomic-dimension meta-analysis was used to investigate dimension-specific contributions to the brain systems underlying CAN activity. To avoid statistical power reduction we capitalized on the covariate approach in ABC. Based on the distribution of the studies, we specified experiments testing the sympathetic system versus other experiments (parasympathetic and the sympathetic/parasympathetic system).

### 2.5. Functional characterization of the clusters of significant ANS activity

A meta-analytic network level characterization was employed to determine whether the identified regions are part of an overarching coactivation involving large scale networks and whether the overarching network resembles more the salience or the default mode networks. To obtain robust results, we performed the following steps separately for ABC and GingerALE.

First, for each identified cluster of CAN activity we computed a single robust condition-independent functional network (Rottschy et al., 2012; Schnellbächer et al., 2020) employing Neurosynth (http://neurosvnth.org) (Yarkoni et al., 2011)). This platform allows the production of seed-based meta-analytic coactivation maps (coactivation maps) and seed-based resting-state functional magnetic resonance imaging (rs-fMRI) maps. The coactivation maps are obtained from the automatic metaanalysis of all fMRI studies present in Neurosynth. These z-score maps (FDR-corrected, (q)=0.01) compute the correlation between the presence/absence of the activity in the defined seed and the presence/absence of the activity observed in each voxel of the brain during the execution of tasks (https://www.neurosynth.org/locations/). These maps, therefore, identify regions that are coactivated with the selected seed during the execution of the fMRI tasks present in the database. The rs-fMRI maps (Pearson’s correlation coefficients map) are obtained from a large dataset of 1000 individuals (provided in Neurosynth as a courtesy of T. Yeo, R. Buckner, and the Brain Genomics Superstructure Project (Buckner et al., 2011; Yeo et al., 2011) and quantify the correlation between the rs-fMRI activity of the seed and the activity observed in each brain vertex (4×4 mm vertex, but to be considered more extended, see (Yeo et al., 2011) for details).

For each identified cluster we built a seed (6 mm spheres centered at the peak coordinates of the clusters transferred in the MNI space), and produced the corresponding coactivation map and rs-fMRI map (thresholded at r>0.2) (Yeo et al., 2011) to identify the connected functional networks during the execution of the tasks (“task-based”) and resting-state conditions (“task-free”), respectively. Then, for each cluster, we defined a single robust functional condition-independent network (from the “task-free” and “task-based” condition), computing the *conjunction map* from the corresponding coactivation map and rs-fMRI map (converted in z-scores), employing the minimum statistics approach (Jakobs et al., 2012). Second, we multiplied the binary images of all the identified *conjunction maps* to assess whether these functional circuits converged in one single network (hereinafter termed *convergent network*).

Third, based on our hypothesis, we next identified whether the *converging network map* was part of the salience network and/or default mode network. Regions of interest of these rs-fMRI networks were obtained from another independent dataset freely available at http://findlab.stanford.edu/research.html (Shirer et al., 2012). To identify the proportion of voxels of each rs-fMRI network comprising the *converging network map*, we multiplied the *converging network map* with the mask of the rs-fMRI map of interest. To strengthen these results, we also computed the proportion of voxels of the rs-fMRI map of interest comprising the *conjunction maps* obtained from each of the identified clusters.

## 3. Results

### 3.1. Study sample characteristic

Based on the inclusion criteria, we identified 42 records (for details see table 1) testing 792 individuals, reporting 44 different experiments (records ID 985 and ID 2645 reported results for two different tasks, respectively somatosensory-motor and task-free, and cognitive and task-free), and 52 different measures of ANS activity (7 studies employed 2 different autonomic measures and specifically: ID 1041, 2470, and 50 employed sympathetic and sympathetic/parasympathetic measures, ID 2403 and ID 2320 parasympathetic and sympathetic/parasympathetic measures, ID 909, ID 515 and 29B two different sympathetic/parasympathetic measures). With respect to the *task dimension*, the affective experiments were the most represented (14 out of 44; 32% of the total amount of experiments) followed by the somatosensory-motor and task-free experiments (both 12 out of 44; 27% of the total experiments), with the least represented experiments being studies employing cognitive tasks (6 out of 44; 14% of the total experiments) (figure 2). With respect to the *autonomic dimension*, experiments testing the sympathetic activity and the sympathetic/parasympathetic activity were equally represented (44%, 23 out of 52 different measures of ANS). Only 12% (6 out of 52 different measures of ANS) examined parasympathetic activity (figure 2).

**Table 1.**
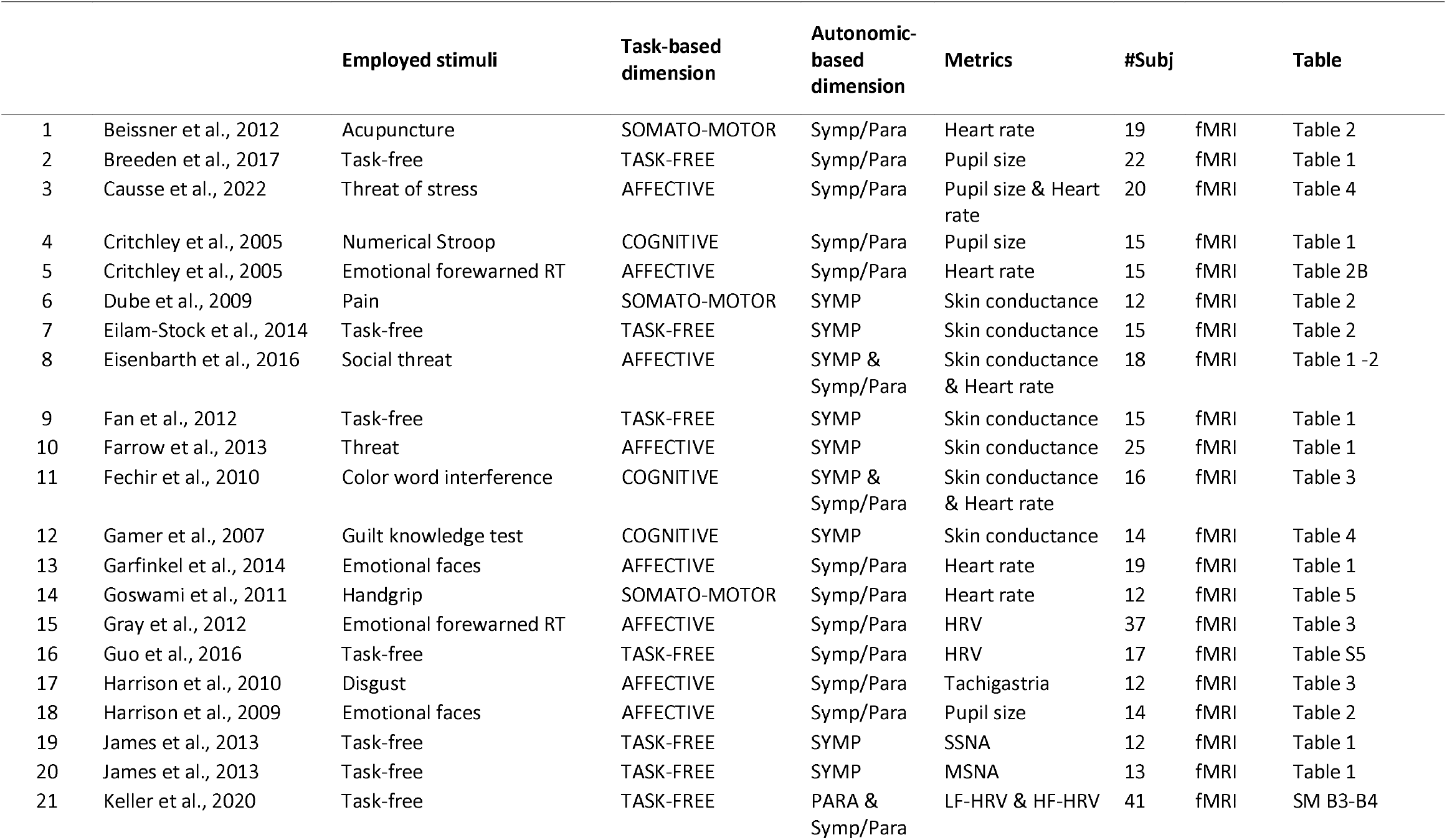
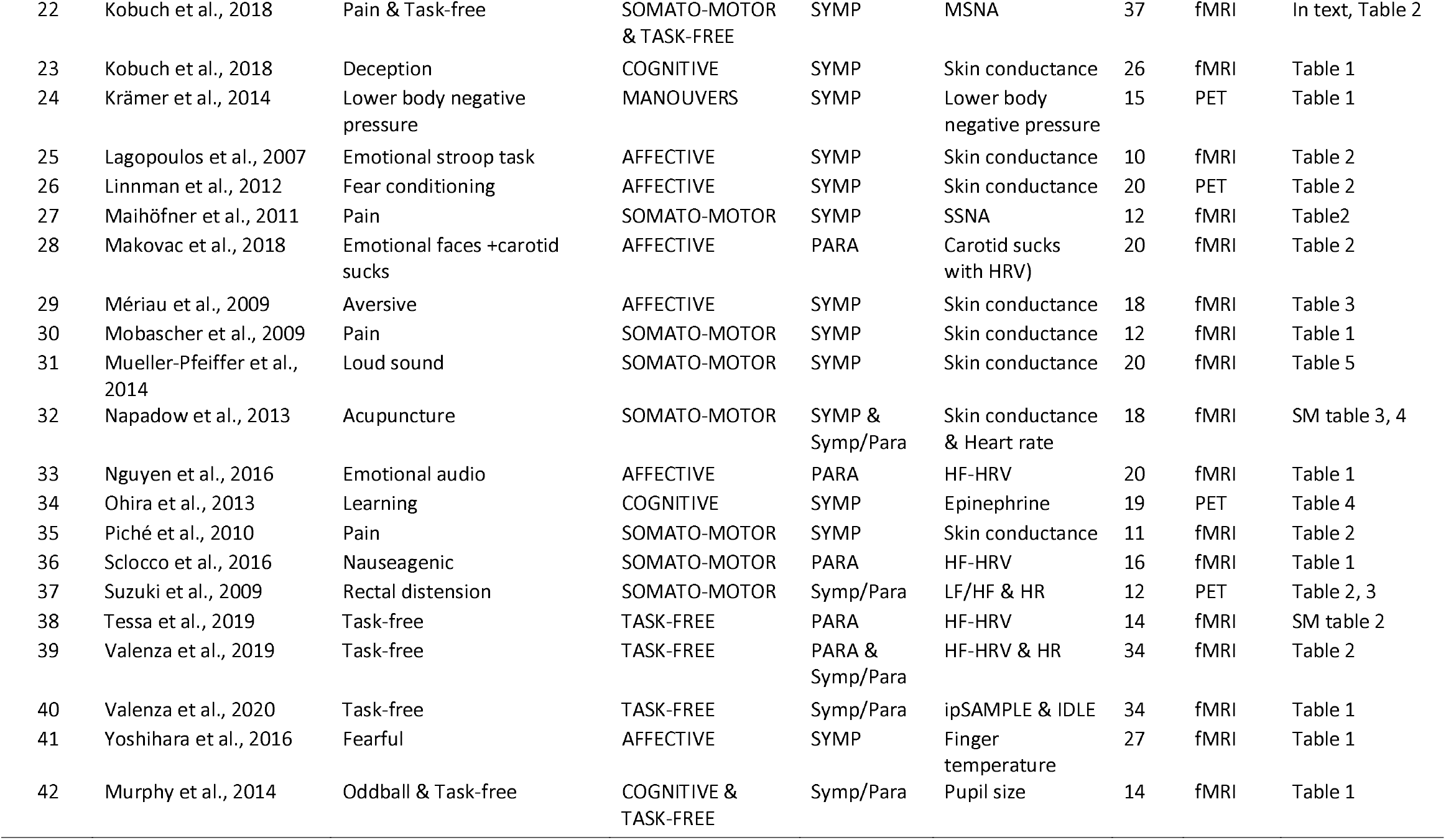
Included studies with their characteristics.

|  |  | Employed stimuli | Task-based dimension | Autonomic-based dimension | Metrics | #Subj |  | Table |
| --- | --- | --- | --- | --- | --- | --- | --- | --- |
| 1 | Beissner et al., 2012 | Acupuncture | SOMATO-MOTOR | Symp/Para | Heart rate | 19 | fMRI | Table 2 |
| 2 | Breeden et al., 2017 | Task-free | TASK-FREE | Symp/Para | Pupil size | 22 | fMRI | Table 1 |
| 3 | Causse et al., 2022 | Threat of stress | AFFECTIVE | Symp/Para | Pupil size & Heart rate | 20 | fMRI | Table 4 |
| 4 | Critchley et al., 2005 | Numerical Stroop | COGNITIVE | Symp/Para | Pupil size | 15 | fMRI | Table 1 |
| 5 | Critchley et al., 2005 | Emotional forewarned RT | AFFECTIVE | Symp/Para | Heart rate | 15 | fMRI | Table 2B |
| 6 | Dube et al., 2009 | Pain | SOMATO-MOTOR | SYMP | Skin conductance | 12 | fMRI | Table 2 |
| 7 | Eilam-Stock et al., 2014 | Task-free | TASK-FREE | SYMP | Skin conductance | 15 | fMRI | Table 2 |
| 8 | Eisenbarth et al., 2016 | Social threat | AFFECTIVE | SYMP & Symp/Para | Skin conductance & Heart rate | 18 | fMRI | Table 1 -2 |
| 9 | Fan et al., 2012 | Task-free | TASK-FREE | SYMP | Skin conductance | 15 | fMRI | Table 1 |
| 10 | Farrow et al., 2013 | Threat | AFFECTIVE | SYMP | Skin conductance | 25 | fMRI | Table 1 |
| 11 | Fechir et al., 2010 | Color word interference | COGNITIVE | SYMP & Symp/Para | Skin conductance & Heart rate | 16 | fMRI | Table 3 |
| 12 | Gamer et al., 2007 | Guilt knowledge test | COGNITIVE | SYMP | Skin conductance | 14 | fMRI | Table 4 |
| 13 | Garfinkel et al., 2014 | Emotional faces | AFFECTIVE | Symp/Para | Heart rate | 19 | fMRI | Table 1 |
| 14 | Goswami et al., 2011 | Handgrip | SOMATO-MOTOR | Symp/Para | Heart rate | 12 | fMRI | Table 5 |
| 15 | Gray et al., 2012 | Emotional forewarned RT | AFFECTIVE | Symp/Para | HRV | 37 | fMRI | Table 3 |
| 16 | Guo et al., 2016 | Task-free | TASK-FREE | Symp/Para | HRV | 17 | fMRI | Table S5 |
| 17 | Harrison et al., 2010 | Disgust | AFFECTIVE | Symp/Para | Tachigastria | 12 | fMRI | Table 3 |
| 18 | Harrison et al., 2009 | Emotional faces | AFFECTIVE | Symp/Para | Pupil size | 14 | fMRI | Table 2 |
| 19 | James et al., 2013 | Task-free | TASK-FREE | SYMP | SSNA | 12 | fMRI | Table 1 |
| 20 | James et al., 2013 | Task-free | TASK-FREE | SYMP | MSNA | 13 | fMRI | Table 1 |
| 21 | Keller et al., 2020 | Task-free | TASK-FREE | PARA & Symp/Para | LF-HRV & HF-HRV | 41 | fMRI | SM B3-B4 |
| 22 | Kobuch et al., 2018 | Pain & Task-free | SOMATO-MOTOR & TASK-FREE | SYMP | MSNA | 37 | fMRI | In text, Table 2 |
| 23 | Kobuch et al., 2018 | Deception | COGNITIVE | SYMP | Skin conductance | 26 | fMRI | Table 1 |
| 24 | Krämer et al., 2014 | Lower body negative pressure | MANOUEVERS | SYMP | Lower body negative pressure | 15 | PET | Table 1 |
| 25 | Lagopoulos et al., 2007 | Emotional stroop task | AFFECTIVE | SYMP | Skin conductance | 10 | fMRI | Table 2 |
| 26 | Linnman et al., 2012 | Fear conditioning | AFFECTIVE | SYMP | Skin conductance | 20 | PET | Table 2 |
| 27 | Maihöfner et al., 2011 | Pain | SOMATO-MOTOR | SYMP | SSNA | 12 | fMRI | Table2 |
| 28 | Makovac et al., 2018 | Emotional faces +carotid sucks | AFFECTIVE | PARA | Carotid sucks with HRV) | 20 | fMRI | Table 2 |
| 29 | Mériaux et al., 2009 | Aversive | AFFECTIVE | SYMP | Skin conductance | 18 | fMRI | Table 3 |
| 30 | Mobascher et al., 2009 | Pain | SOMATO-MOTOR | SYMP | Skin conductance | 12 | fMRI | Table 1 |
| 31 | Mueller-Pfeiffer et al., 2014 | Loud sound | SOMATO-MOTOR | SYMP | Skin conductance | 20 | fMRI | Table 5 |
| 32 | Napadow et al., 2013 | Acupuncture | SOMATO-MOTOR | SYMP & Symp/Para | Skin conductance & Heart rate | 18 | fMRI | SM table 3, 4 |
| 33 | Nguyen et al., 2016 | Emotional audio | AFFECTIVE | PARA | HF-HRV | 20 | fMRI | Table 1 |
| 34 | Ohira et al., 2013 | Learning | COGNITIVE | SYMP | Epinephrine | 19 | PET | Table 4 |
| 35 | Piché et al., 2010 | Pain | SOMATO-MOTOR | SYMP | Skin conductance | 11 | fMRI | Table 2 |
| 36 | Sclocco et al., 2016 | Nauseagenic | SOMATO-MOTOR | PARA | HF-HRV | 16 | fMRI | Table 1 |
| 37 | Suzuki et al., 2009 | Rectal distension | SOMATO-MOTOR | Symp/Para | LF/HF & HR | 12 | PET | Table 2, 3 |
| 38 | Tessa et al., 2019 | Task-free | TASK-FREE | PARA | HF-HRV | 14 | fMRI | SM table 2 |
| 39 | Valenza et al., 2019 | Task-free | TASK-FREE | PARA & Symp/Para | HF-HRV & HR | 34 | fMRI | Table 2 |
| 40 | Valenza et al., 2020 | Task-free | TASK-FREE | Symp/Para | ipSAMPLE & IDLE | 34 | fMRI | Table 1 |
| 41 | Yoshihara et al., 2016 | Fearful | AFFECTIVE | SYMP | Finger temperature | 27 | fMRI | Table 1 |
| 42 | Murphy et al., 2014 | Oddball & Task-free | COGNITIVE & TASK-FREE | Symp/Para | Pupil size | 14 | fMRI | Table 1 |

**Fig. 2.**
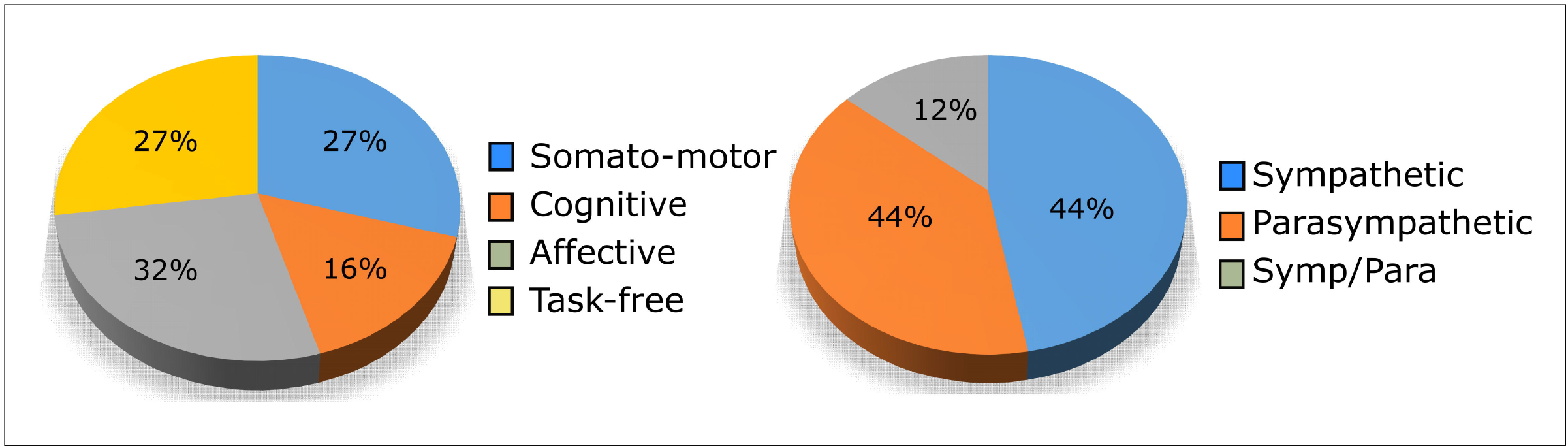
Graphical representations of the percentages of the selected studies according to their taskdimension (on the left) and autonomic dimension (on the right). Abbreviations: Somato-motor: somatosensory-motor experiments; Symp: sympathetic experiments; Para: parasympathetic experiments; Symp/Para: sympathetic and parasympathetic experiments.

In line with the approach by Beissner et al. (Beissner et al., 2013) the selected studies employed a dimensional regression approach identifying regions that linearly increased their activity as a function of the ANS index as well as categorical comparisons of the levels of ANS activity between different conditions. Capitalizing on both approaches, allowed us to increase statistical power and determine ANS irrespective of the employed analysis.

### 3.2. Meta-analytic results

#### Central autonomic network – pooled meta-analyses

Across all selected experiments, ABC and GingerALE presented very similar meta-analytic results (table 2, figure 3). ABC revealed three significant clusters located in the right midcingulate cortex (MNI peak coordinates in [3 9 43], 12 contributing experiments), left dorsal anterior insula cortex (MNI peak coordinates in [-38 5 142], 6 contributing experiments), and right dorsal anterior insula cortex (peak coordinates in [40 19 −2], 8 contributing experiments) (figure 4). Very similarly, GingerALE revealed three clusters with their peaks located in the right midcingulate cortex (MNI peak coordinates in [4 8 44], 14 contributing experiments), and in the dorsal anterior section of the left insula (MNI peak coordinates in [-38 5 11], 9 contributing experiments), and of the right insula (peak coordinates in [34 20 4], 18 contributing experiments). According to a probabilistic atlas of the insula cortex (Faillenot et al., 2017), the left insula cluster encompassed the dorsal regions of the middle and posterior short gyrus, while the right cluster was located on the dorsal region of the anterior and middle short gyrus (figure 3), for both ABC and GingerALE algorithms.

**Table 2.**
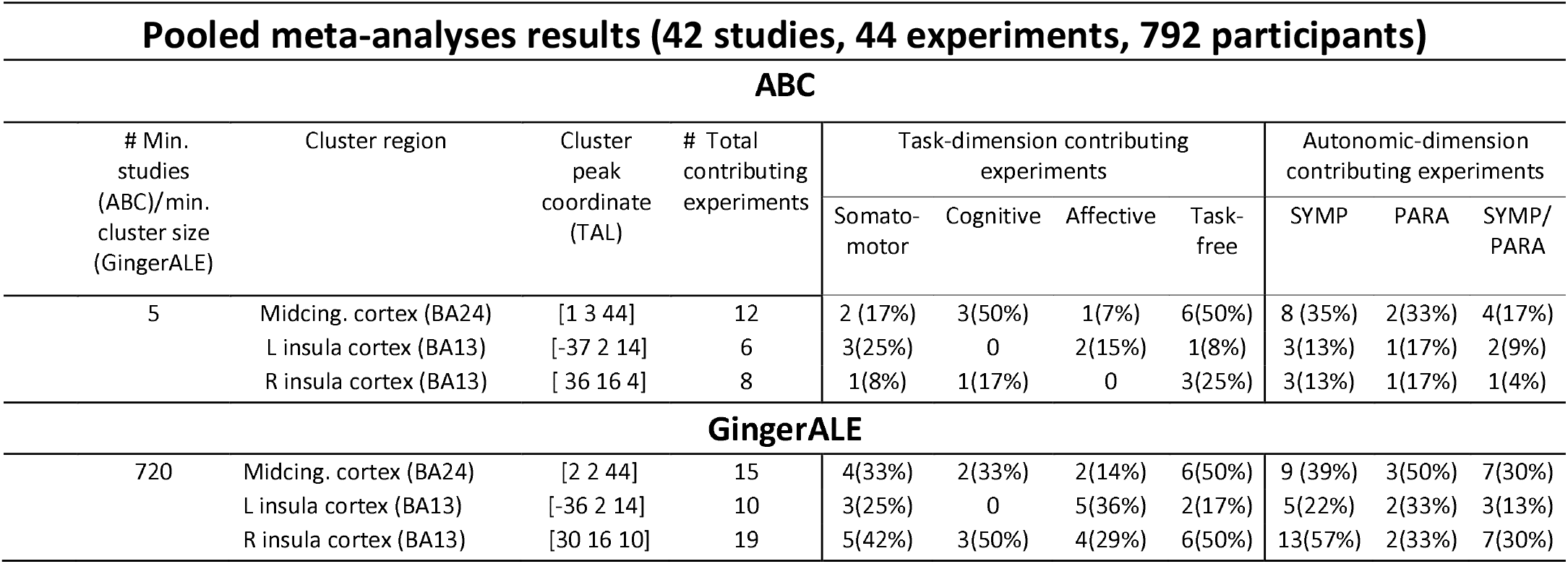
Pooled meta-analyses result for ABC and GingerALE. R = right; L = left

**Fig. 3.**
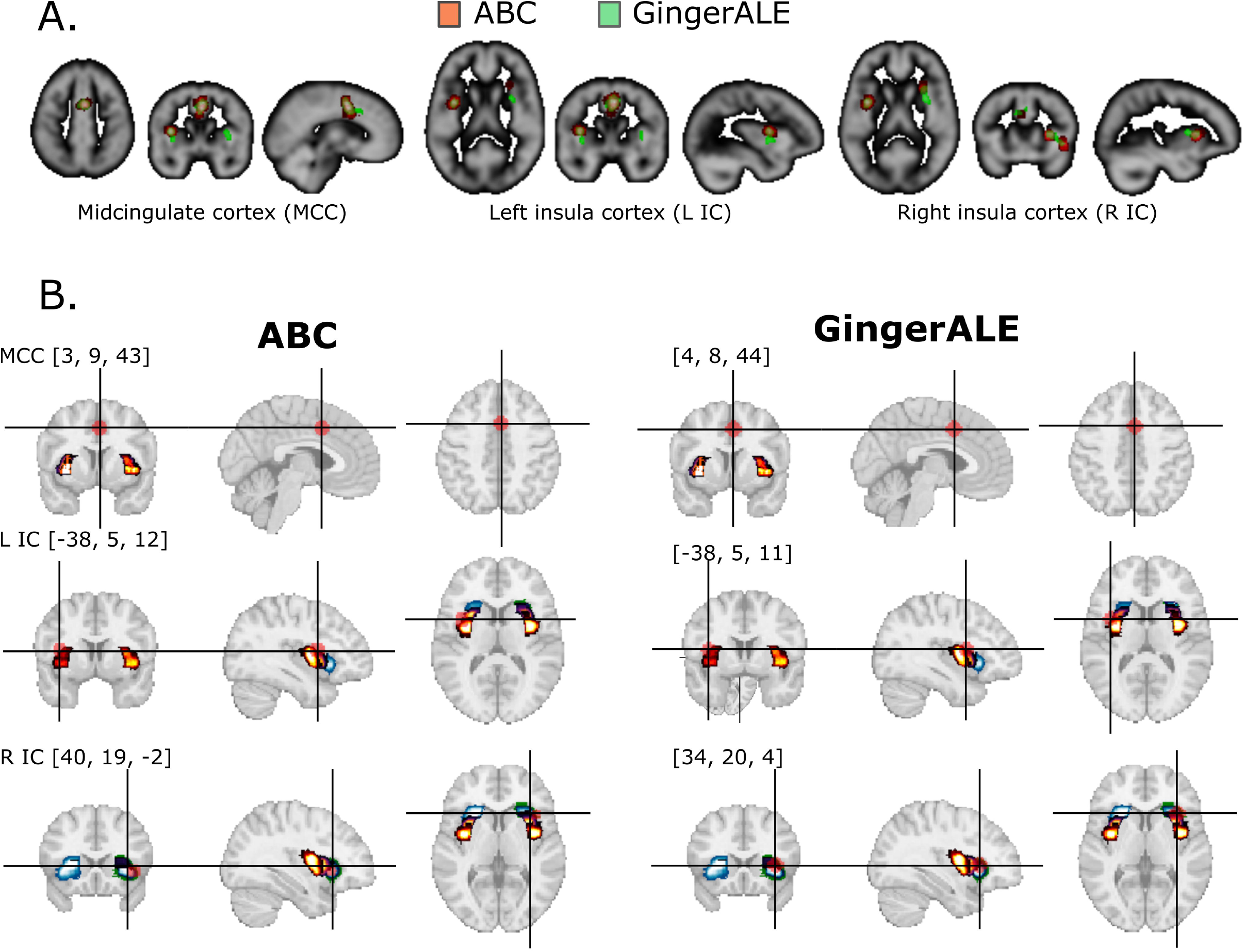
**A.** Representations in the TAL space of the identified clusters for ABC (in red) and GingerALE (in green) for the pooled meta-analysis. **B.** Position of each seed computed as 6mm sphere centered in the peak coordinates of the identified meta-analytic clusters (in midcingulate cortex, left insula cortex, and right insula cortex) for ABC and GingerALE. These seeds were used to produce meta-analytic coactivation maps and rs-fMRI maps employing Neurosynth (Yarkoni et al. 2011).

**Fig. 4.**
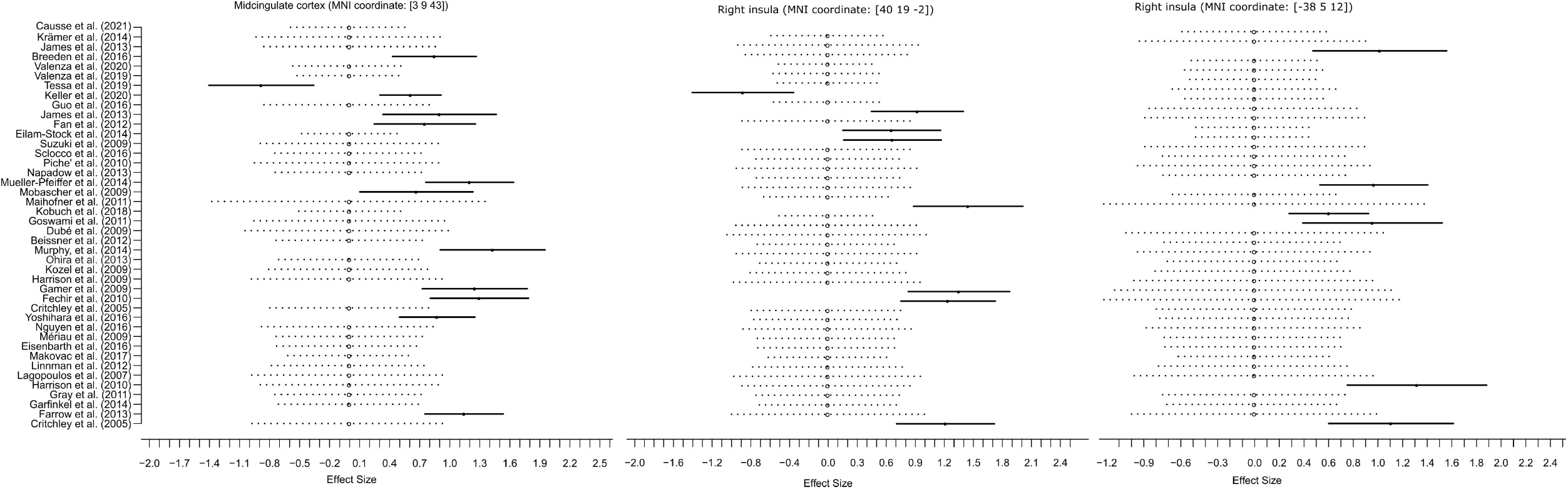
Forest plots for each identified cluster showing the effect estimates and confidence intervals of the individual studies in the meta-analysis computed with ABC (Tench et al. 2022).

#### Task-dimension meta-analyses

The ABC meta-analysis employing *task dimension* as covariate of interest revealed a significant effect of task-free and cognitive experiments versus somatosensory-motor and affective experiments on the cluster identified in the midcingulate cortex (binary logistic regression: 95% C.I.: 0.313 - 3.487, OR = 6.685, p-value=0.016) (table 3). This cluster emerged from the convergent activity of 6 task-free (50% of task-free experiments) and 3 cognitive (43% of cognitive experiments) experiments, and only of 2 somatosensory-motor (17% of somatosensory-motor experiments) and 1 affective (7% of affective) experiments.

**Table 3.** Logistic regression model for the task-dimension (task-free and cognitive experiments vs. somatosensory-motor and affective experiments), and the autonomic dimension (sympathetic vs. parasympathetic and the sympathetic/parasympathetic system) obtained employing ABC (Tench et al. 2022). Midcing. = midcingulate cortex, CI = confidence; OR = odds ratio, R = right; L = left.

| ABC - pooled meta-analysis with covariates |  |  |  |  |  |  |  |  |  |  |  |
| --- | --- | --- | --- | --- | --- | --- | --- | --- | --- | --- | --- |
|  |  | Logistic regression model for<br>'Task-dimension' |  |  |  |  | Logistic regression model for<br>'Autonomic -dimension' |  |  |  |  |
|  |  | Parameter | CI |  | OR | P-value | Parameter | CI |  | OR | P-value |
| Midcing cortex (BA24) | [2 4 44] | 1.90 | 0.31 | 3.49 | 6.69 | 0.02 | 0.92 | -0.51 | 2.36 | 2.52 | 0.20 |
| L insula cortex (BA13) | [-36 2 14] | -0.15 | -2.07 | 1.77 | 0.86 | 0.87 | 0.54 | -1.51 | 2.58 | 1.71 | 0.60 |
| R insula cortex (BA13) | [ 44 16 0] | 1.83 | 0.00 | 3.66 | 6.23 | 0.05 | 0.30 | -1.51 | 2.10 | 1.35 | 0.74 |

#### Autonomic-dimension meta-analyses

The ABC meta-analysis employing *autonomic dimension* as covariate of interest revealed that experiments examining the sympathetic system did not contribute differently to the identified clusters when compared to the remaining experiments pooling studies examining parasympathetic and sympathetic/parasympathetic system activity This indicates that all the studies contributed equally to the activity in the observed clusters independently from the autonomic system involved (see table 3).

### 3.3. Functional characterization

#### Pooled analyses

The *conjunction maps* (table 4) obtained for each identified cluster from the resting state functional connectivity (rs-FC) map and the meta-analytic co-activation map, revealed highly similar networks for both ABC and GingerALE meta-analyses (see figure 5). The computation of the proportion of *conjunction map* voxels overlapping the specific rs-fMRI network showed that the identified networks mainly constitute the salience network according to the rs-fMRI dataset employed (ABC: from 34% to 50% of the salience network voxels was part of each single conjunction map; GingerALE: from 34% to 56%). In contrast, no evidence was found that these conjunction maps were part of the default mode network (maximum 1% of the voxels of the default mode network comprised these conjunction maps).

**Table 4.**
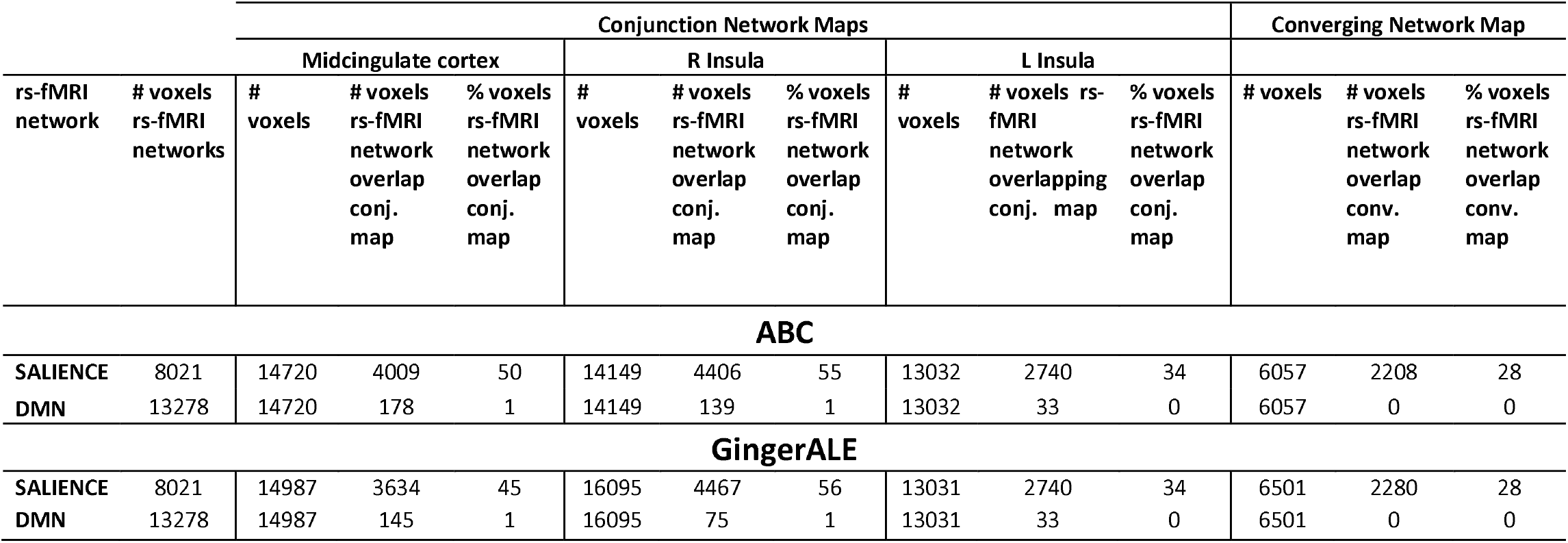
Results of the functional characterizations. Number of total voxels of the employed rs-fMRI networks (salience network and default mode network), number of total voxels of the conjunction network map built for each identified cluster, number of total voxels of the converging network map, and number of voxels and relative percentages of the considered rs-fMRI networks overlapping the conjunction maps and of the converging network map for ABC (Tench et al. 2022) and GingerALE (Eickoff 2009,2012). Conj = conjunction map; Con = convergent map, DMN = default mode network.

|  |  | Conjunction Network Maps |  |  |  |  |  |  |  |  | Converging Network Map |  |  |
| --- | --- | --- | --- | --- | --- | --- | --- | --- | --- | --- | --- | --- | --- |
|  |  | Midcingulate cortex |  |  | R Insula |  |  | L Insula |  |  |  |  |  |
| rs-fMRI network | # voxels rs-fMRI networks | # voxels | # voxels rs-fMRI network overlap conj. map | % voxels rs-fMRI network overlap conj. map | # voxels | # voxels rs-fMRI network overlap conj. map | % voxels rs-fMRI network overlap conj. map | # voxels | # voxels rs-fMRI network overlapping conj. map | % voxels rs-fMRI network overlap conj. map | # voxels | # voxels rs-fMRI network overlap conv. map | % voxels rs-fMRI network overlap conv. map |
| ABC |  |  |  |  |  |  |  |  |  |  |  |  |  |
| SALIENCE | 8021 | 14720 | 4009 | 50 | 14149 | 4406 | 55 | 13032 | 2740 | 34 | 6057 | 2208 | 28 |
| DMN | 13278 | 14720 | 178 | 1 | 14149 | 139 | 1 | 13032 | 33 | 0 | 6057 | 0 | 0 |
| GingerALE |  |  |  |  |  |  |  |  |  |  |  |  |  |
| SALIENCE | 8021 | 14987 | 3634 | 45 | 16095 | 4467 | 56 | 13031 | 2740 | 34 | 6501 | 2280 | 28 |
| DMN | 13278 | 14987 | 145 | 1 | 16095 | 75 | 1 | 13031 | 33 | 0 | 6501 | 0 | 0 |

**Fig. 5.**
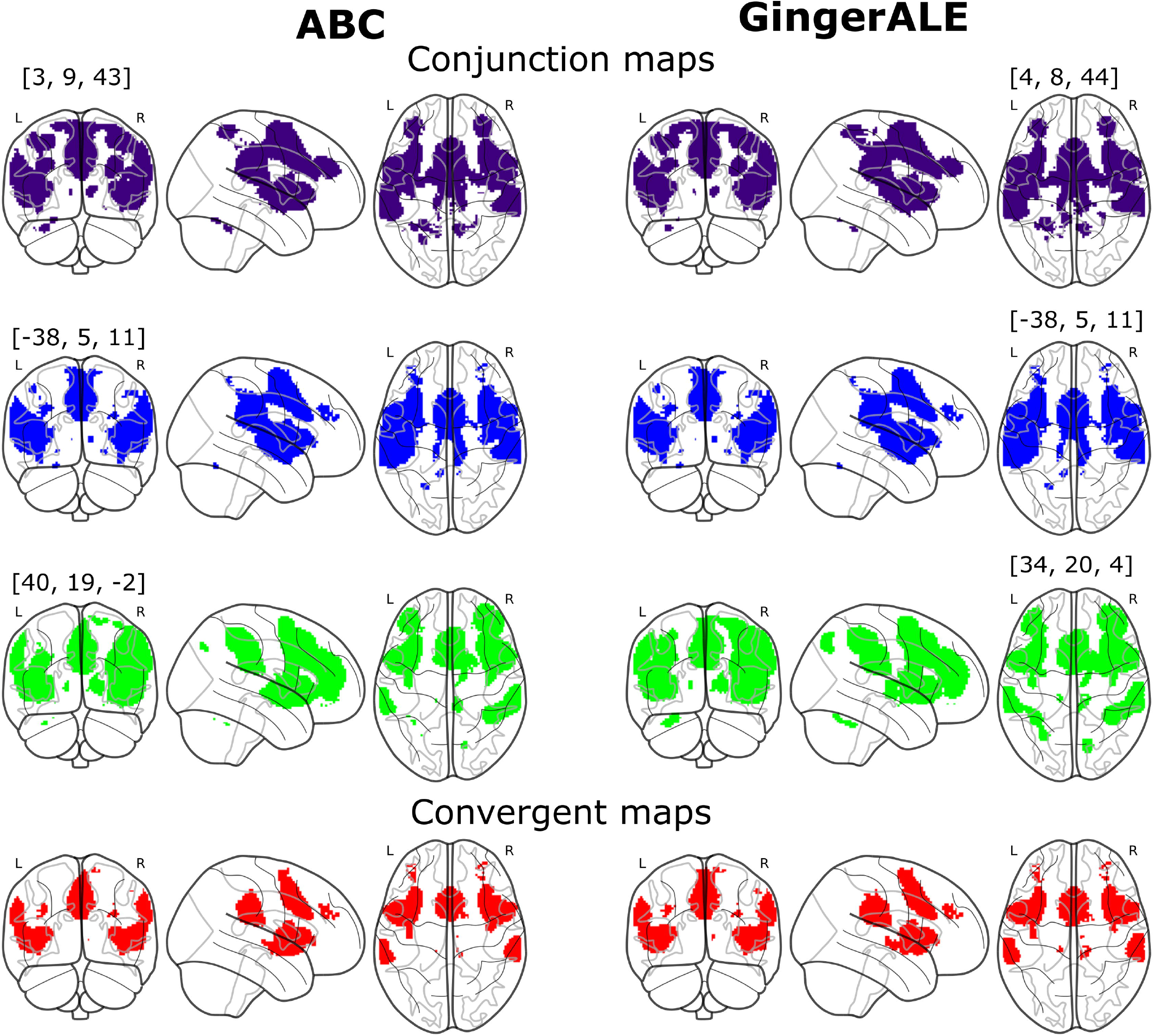
Glass-brain representations: Conjunctions maps computed employing the minimum statistical approach between the meta-analytic coactivation map and the rs-fMRI map obtained from each seed computed as 6mm sphere centered in the peak coordinates of the identified meta-analytic clusters (in midcingulate cortex, left insula cortex, and right insula cortex) and the corresponding Convergent network maps for ABC and GingerALE.

Importantly, computation of the *converging network map* (table 5) for each meta-analytic tool showed that the conjunction maps of each identified cluster converged in one single map comprising the bilateral anterior/middle insula cortex and the midcingulate cortex, but also the bilateral inferior parietal lobule and small clusters in the bilateral middle frontal gyrus (see figure 5, 6A and 6B). Also in this case, a good proportion of the salience network contained the *converging network* (ABC and GingerALE: 28% of the salience network). As expected, based on the results obtained from the *conjunction maps*, no evidence was found that the default mode network was contained in the converging network (0% of the voxels of the default mode network comprised the *converging network* (see figure 6A and 6B and table 4).

**Fig. 6.**
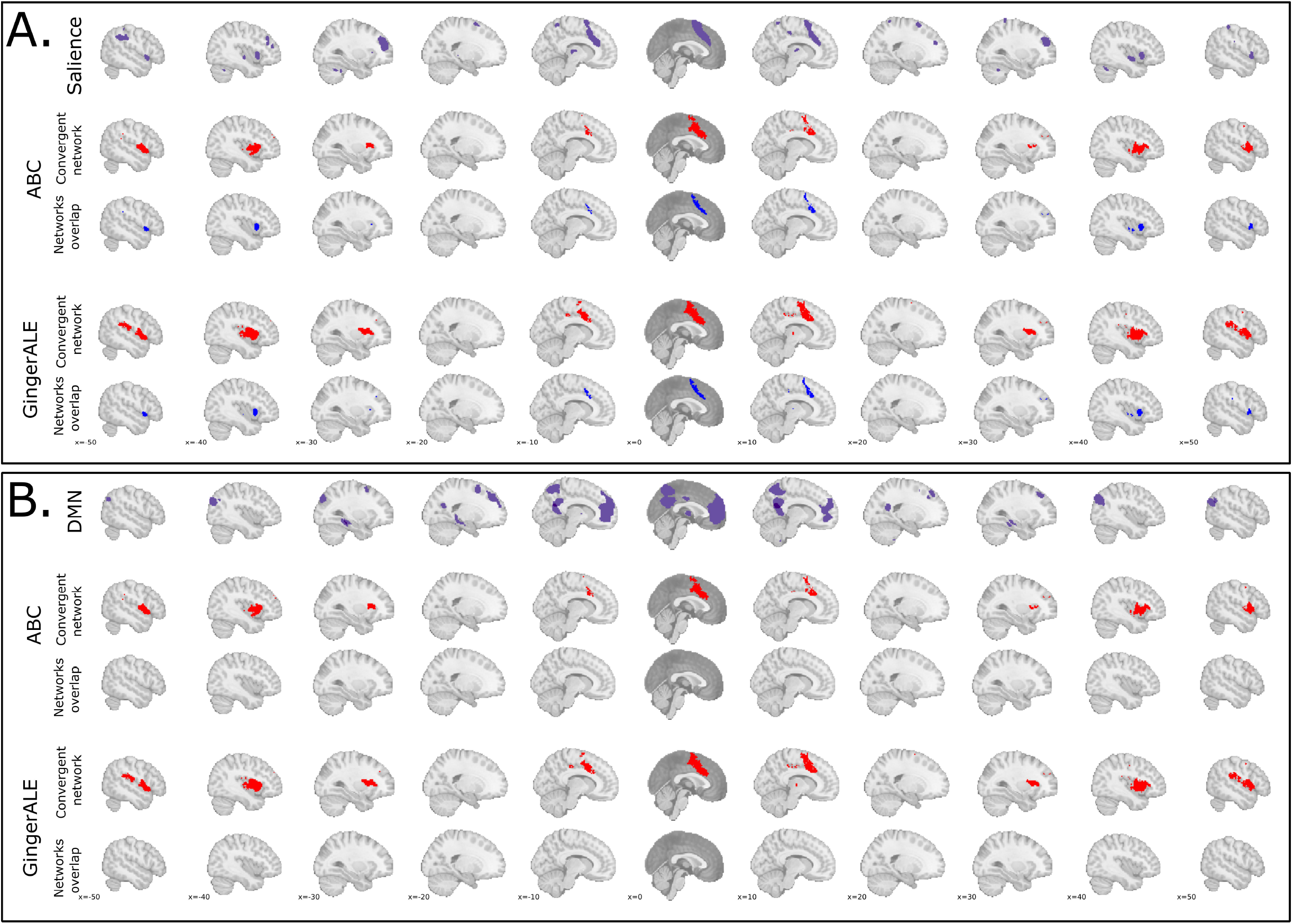
A. Graphical representations of the Salience network map obtained from an independent dataset (http://findlab.Stanford.edu/functional\_ROIs.html), of the Convergent network map and of the results of the overlap between the two (Networks overlap map). B. Graphical representations of the Default mode network map obtained from an independent dataset (http://findlab.Stanford.edu/functional\_ROIs.html), of the Convergent network map and the results of the overlap between the two (Networks overlap map).

**Table 5.** Cortical regions (extracted with Anatomy toolbox) of the convergent network maps obtained from the clusters identified with ABC (Tench et al. 2022) and GingerALE (Eickoff 2009,2012). R = right; L = left.

| Clusters | n of voxels | Center of gravity (MNI coordinates) | Identified Brain regions |
| --- | --- | --- | --- |
| **ABC** |  |  |  |
| **cluster #1** | 1771 | [44 10 4] | R Insula |
| **cluster #2** | 1771 | [-42 8 4] | L Insula |
| **cluster #3** | 1439 | [1 13 40] | L & R Midcingulate cortex |
| **cluster #4** | 648 | [62 -20 30] | R Inferior parietal lobule |
| **cluster #5** | 466 | [-61 -32 27] | L Inferior parietal lobule |
| **GingerALE** |  |  |  |
| **cluster #1** | 1684 | [44 10 4] | R Insula |
| **cluster #2** | 1674 | [1 11 42] | L & R Midcingulate cortex |
| **cluster #3** | 1582 | [-42 8 5] | L Insula |
| **cluster #4** | 786 | [61 -27 29] | R Inferior parietal lobule |
| **cluster #5** | 622 | [-60 -30 27] | L Inferior parietal lobule |

## 4. Discussion

Capitalizing on recent advances in CBMA and two different algorithms (ABC and GingerALE) in combination with an increasing number of recent human neuroimaging studies we determined insular and cingulate areas of meta-analytic convergence in human forebrain CAN. Furthermore, we identified the possible underlying functional network involved in ANS processing and modulation using an unbiased approach. In the context of the goals of the study three robust findings were observed. First, across both CBMA algorithms the bilateral dorsal anterior insula and midcingulate cortex were robustly engaged in CAN. Second, while the bilateral dorsal anterior insula proved to be involved in processing ANS signals across both affective and non-affective task domains (i.e., somatosensory-motor and affective task vs task-free and cognitive task domain), the midcingulate cortex appeared to be primarily involved in processing ANS signals during cognitive tasks and task-free conditions. Third, the functional connectivity networks determined for each identified cluster engaged a single functional condition-independent integrated circuit located partially in the salience network. The analyses additionally determined the bilateral inferior parietal lobules as nodes of this functional network, however no evidence for an involvement of the default mode network was found.

Our finding that the bilateral dorsal anterior insular cortex is involved in the modulation of both autonomic systems during all task types supports the hypothesis that these regions are fundamental hubs of CAN. This finding, coupled with the evidence that the functional network supporting CAN is at least partially anchored in the salience network is supported by evidence from different lines of research such that a long tradition of animal and human studies has shown the anterior insular cortex is a critical viscerosensory area, being the primary site of cortical interoceptive projection and strongly involved in interoceptive processing and interoceptive dysregulations (Arthur D Craig, 2003; Arthur D Craig & Craig, 2009b; Critchley et al., 2013; Ferraro et al., 2021; Foxe & Schroeder, 2005; Li et al., 2018; Uddin, 2014; Xu et al., 2020; Xue Hai, 2016). One of the primary functions of this region, and aligning with its involvement in a wide range of conditions and behaviors (Arthur D Craig & Craig, 2009a; Uddin, 2014), appears to be the integration of external information with the internal state of the body (interoception) and the prioritization of stimuli through the flow of internal and external information based on their salience to the individual (Menon, 2011; Uddin, 2014; Yao et al., 2018). Consistent with this hypothesis, the anterior insular cortex, as a key hub of the ventral salience and attention system (Fox & Raichle, 2007; Menon, 2011; Seeley et al., 2007a; Uddin, 2014), has been shown to control the switch between the activity of the default mode network, implicated in self-referential operations, and the dorsal attention network, dedicated to top-down attentional resource allocation (Huang et al., 2021; Xin et al., 2021). Recent work has also provided strong evidence that this region may be critical for the conscious experience not only of pain (Bastuji et al., 2018) but also of sensory information (Huang et al., 2021), with heart-insula interactions appearing to play a critical role in body self-identification (Park et al., 2018). Our results thus support the hypothesis that the bilateral dorsal anterior insula is the fundamental cortical hub involved in regulating both autonomic systems regardless of negative affect.

Our results additionally revealed another cortical region exhibiting meta-analytic convergence involved in autonomic modulations: the midcingulate cortex. In the context of autonomic modulations, the midcingulate cortex has been implicated in a wide range of cognitive functions (Etkin et al., 2011; Heilbronner & Hayden, 2016; Kober et al., 2008; Vogt, 2016), prepares individuals for action through modulation of internal body state (Beissner et al., 2013; Touroutoglou et al., 2020) and integrates negative affect and cognition (Shackman et al., 2011; Tolomeo et al., 2016). Altough the midcingulate was identified together with the dorsal insula it differed with respect to the negative affective load of the task domain. We observed that the midcingulate cortex activity is strongly dependent on the type of ongoing task, being mainly supported by cognitive manipulations and task-free conditions, where the experienced negative affect is, in principle, relatively lower in comparison to the negative affect experienced during affective (mainly inducing negative emotions) or somatosensory-motor tasks (mainly inducing acute pain). Importantly, this result was not a bias of the different autonomic systems investigated, as indicated by the Freeman-Halton extension of Fisher’s exact probability test showing that the type of employed task and the investigated autonomic system are independent variables. The midcingulate cortex plays an important role in linking interoceptive signals with cognitive and negative affective processes (Shackman et al., 2011; Tolomeo et al., 2016) and in the modulation of body-arousal states (Critchley et al., 2003) by demands posed by the environment. In the present study the midcingulate cortex appears to be involved in the modulation of the autonomic system depending on the negative affective component of the context, possibly reflecting the rather ‘active state’ during the negative affect-free ANS engagement (i.e., cognitive tasks or task-free conditions characterized by an intense and not constrained self-referential activity) while in contrast modulation of the autonomic system may rather occur during negative affect induction and a corresponding ‘passive state’. This result supports the hypothesis that CAN could have a highly dynamic organization with involvement of the midcingulate cortex primarily related to its role as an interface between interoception and action as a visceral motor cortex (Saper & Stornetta, 2015).

The functional characterization of the identified clusters (i.e., the bilateral anterior insula cortex and middle cingulate cortex) showed that these regions were part of a single symmetrical and bilaterally distributed functional network that includes, in addition to the three main identified hubs, two symmetrical areas of the bilateral inferior parietal lobule and small clusters of the middle frontal gyrus. It is worth noting that this result was obtained by employing the seed-based rs-fMRI maps of a large publicly available dataset and the seed-based meta-analytic fMRI coactivation maps produced by the Neurosynth platform. The identified condition-independent network overlapped considerably with the salience network and appears to constitute a functional circuit underlying the central processing of ANS signals. The inferior parietal lobule plays a decisive role in the representation of own body (Berlucchi & Aglioti, 1997), and along with the insular cortex and frontal lobes is involved in the construction of body awareness and sense of self (Berlucchi & Aglioti, 1997, 2010; Melzack, 1990). These multidimensional constructs have their foundation in interoceptive signals (A. D. Craig, 2002; Critchley & Harrison, 2013) arising from ANS activity (Berlucchi & Aglioti, 1997; Blanke et al., 2015; Tanaka et al., 2021). Along these lines, the bilateral inferior parietal lobule has been shown to be the site of integration of interoceptive and exteroceptive information (Salvato et al., 2019). These findings substantiate the claim that the bilateral anterior insula cortex, the midcingulate cortex, and the bilateral inferior parietal lobule, with small regions of the bilateral middle frontal gyrus, constitute a single robust functional network supporting a central representation of the autonomic systems. Inconsistent with assumptions of previous work, our results do not provide meta-analytic support either for the presence of divergent networks underlying sympathetic and parasympathetic systems (Beissner et al., 2013) or for a robust involvement of the default mode network in central processing of autonomic activity (Beissner et al., 2013; Nagai et al., 2004; Thayer et al., 2012), in particular parasympathetic activity (Babo-Rebelo et al., 2016; Ruffle et al., 2021). These results, however, should be taken with caution because they may be due to the effects of the relatively low number of studies investigating the parasympathetic system (12%) compared with studies investigating the sympathetic system (44%) and the sympathetic/parasympathetic system (44%), making our results more consistent with the central processing network of sympathetic activity. However, it is important to note that 3 of the 6 studies investigating central processing of parasympathetic activity helped identify meta-analytic clusters observed in the midcingulate cortex and left anterior/middle insular cortex, suggesting that central processing of the sympathetic and parasympathetic systems occurs along the same brain pathways.

The present meta-analysis represents an update to the seminal meta-analysis examining the neural basis of the CAN nearly 10 years ago by Beissner and colleagues (Beissner et al., 2013) and capitalized on an increasing number of studies as well as conceptual and methodological progress. Despite the convergent results regarding g the role of the insula and midcingulate some differences emerged. The CAN reported in Beissner et al. (Beissner et al., 2013) encompassed several cortical areas (anterior and middle cingulate cortex, ventral posterior cingulate cortex, bilateral anterior insula, right frontal insular cortex left posterior insular cortex, and ventromedial prefrontal cortex, and lateral parietal area - including the right angular gyrus and supramarginal gyrus) as well as subcortical structures (thalamus, bilateral amygdala, right hippocampal formation, hypothalamus, midbrain, and brainstem regions) and showed that sympathetic and parasympathetic modulations map to different areas while the DMN appeared to be involved in parasympathetic system. In contrast to the widespread cortical and subcortical areas of CAN observed by Beissner et al. (Beissner et al., 2013), our meta-analysis provided evidence of only three clusters located respectively in the bilateral dorsal anterior insular and the midcingulate cortex (partially overlapping with the regions identified by Beissner et al. (Beissner et al., 2013) as central regulatory areas of autonomic activity). Furthermore, contrary to findings in (Beissner et al., 2013), our study showed that the midcingulate cortex would regulate the autonomic system primarily during cognitive and task-free experiments, whereas activity in the bilateral dorsal anterior insular cortex would modulate ANS activity regardless of task type. As final points of inconsistency, we found no divergent activity between the sympathetic and parasympathetic systems either in terms of laterality or in terms of areas, nor did we find involvement of the DMN as a regulatory network of the parasympathetic activity, nor the involvement of the left amygdala as a crucial modulatory component of both autonomic systems.

Several factors might have led to these inconsistencies. First, it is well recognized that to obtain a robust meta-analysis, there is a need for investigations of high methodological quality that can produce a high level of evidence; otherwise, the risk is to have low-quality meta-analyses or even erroneous results (Guyatt et al., 2008). In this regard, the neuroimaging community still lacks standardized tools that can define the level of methodological quality for studies to be selected in a meta-analysis. We, therefore, sought to reduce the possible bias induced by papers characterized by low statistical power by excluding studies that investigated CAN in fewer than 10 individuals. Given the evolution of methodological aspects in neuroimaging research, most of these studies were published before 2010, drastically decreasing the number of common studies between our meta-analysis and that of Beissner et al. (2013). Second, in addition to having selected studies published in recent years, we enriched our pool of selected studies with task-free experiments. Third, the version of the algorithm (GingerALE 2.3) (Eickhoff et al., 2009; Rottschy et al., 2012) employed by Beissner et al. (Beissner et al., 2013) suffered from implementation errors, which were fixed in version 2.3.3 (Eickhoff et al., 2017).

From a global perspective, the main goal of a meta-analysis is to synthesize the available data by generalizing the results of a more significant number of studies, thus providing a complete picture (Gurevitch et al., 2018) with the highest available level of evidence for the topic under investigation. However, in addition to the problems that every meta-analysis brings with it (Gurevitch et al., 2018), meta-analyses on neuroimaging studies adds other levels of complexity: such as the already discussed lack of a standardized assessment of the methodological validity of the selected papers, or the fact that many studies, employing regions of interest analyses, cannot be selected in CBMA since they violate one of its fundamental assumptions, namely that the coordinates of the peaks reported in the studies are uniformly distributed in the gray matter (Turkeltaub et al., 2002). In this context, the case of the amygdala is paradigmatic: despite the abundant literature reporting its involvement in CAN, our meta-analysis was unable to detect this structure as a critical region of this circuit possibly because many neuroimaging studies use only region-of-interest approaches.

Another important region that does not emerge from this meta-analysis is the posterior insular cortex. Primary interoceptive signals are represented in the posterior regions of the insula cortex, while their integration into perceptual maps occurs in the middle and anterior parts of the insula cortex according to a posterior-anterior gradient (Craig 2009). Because of the different autonomic signals used by the various selected studies, our meta-analysis could reveal only the common site of integration of these signals, without identifying the source of such heterogeneous primary interoceptive signals.

In conclusion, our robust meta-analysis suggests that both sympathetic and parasympathetic systems are rooted in a single functional networkpartially overlapping with the salience network and anchored in the bilateral dorsal anterior insular cortex, the midcingulate cortex areas and bilateral inferior parietal lobule. However, while the dorsal anterior insular regions constitute a fundamental hub of CAN representing the interoceptive flow and its integration with external information to guide behavior (Menon, 2011; Uddin, 2014), the midcingulate cortex appears to integrate interoception, cognition, and action, perhaps mobilizing the metabolic resources required to perform engaging tasks through its direct connection to brainstem arousal nuclei (An et al., 1998). This latter evidence, by showing that ANS modulations in a specific CAN region support specific behavior, enriches the notion that ANS is not a monolithic system (Iacovella et al., 2018; Norman et al., 2014),but possibly a hierarchically organized network as postulated by the neurovisceral integration model (Smith et al., 2017; Thayer et al., 2012). Furthermore, the observation that the bilateral inferior parietal lobule is part of the CAN functional network, provides support for the hypothesis that ANS peripheral signals with interoception are the underpinning body awareness and sense of self.

Remarkably, the critical regions of CAN observed in this meta-analysis are among the most reported co-activated areas in neuroimaging studies and have been repeatedly shown to be dysregulated across different mental and neurological disorders (e.g., Ferraro et al., 2021; Goodkind et al., 2015; Klugah-Brown et al., 2021) This suggests that CAN and the dynamic interaction to maintain bodily homeostasis may be present in several brain imaging studies associated with increased ANS engagement and that dysfunction (or disruption) in this interplay may underpin unspecific pathological symptoms across mental and neurological disorders.

## Data availability statement

The selected studies are reported in the manuscript. The coordinates will be provided upon request.

